# Individual variability in sensorimotor learning reflects trait-like neurobehavioral subject factors

**DOI:** 10.1101/2024.04.11.589135

**Authors:** Corson N. Areshenkoff, Anouk J. de Brouwer, Daniel J. Gale, Dominic Standage, Joseph Y. Nashed, J. Randall Flanagan, Jason P. Gallivan

**Affiliations:** Centre for Neuroscience Studies, Queens University, Kingston Ontario; Department of Psychology, Queens University, Kingston Ontario; Department of Biomedical and Molecular Sciences, Queens University, Kingston Ontario

## Abstract

Models of Human motor behaviour often emphasize the computations performed by the motor system during learning. Yet, there is an emerging consensus that the learning of even simple motor actions can be augmented by sophisticated cognitive strategies, which rely upon executive functions implemented throughout the cortex. These executive functions, in turn, have been linked to stable subject differences in intrinsic brain organization function, observable even during rest. Here we show, using behavioural studies in humans, that individual differences in the rate of sensorimotor adaptation are linked to differences in executive function, as assessed using the classic trail-making task. Secondly, using separate train and test functional MRI datasets, we show that specific patterns of resting-state functional connectivity between higher-order cognitive brain networks, which have been previously linked to executive function, subsequently predict more rapid learning during sensorimotor adaptation. Importantly, this relationship was unique to cognitive brain networks, as the functional connectivity between sensorimotor brain networks did not predict subsequent motor learning performance. Together, these findings suggest that individual differences in motor learning reflect, to a substantial degree, trait-like subject differences in cognitive brain network structure. This perspective invites broader consideration as to the origins of motor learning ability, and links motor performance to an expansive literature implicating the coordinated functioning of the default mode, ventral attention, and frontoparietal networks in flexible behavioral control.

---

The human brain possesses a remarkable capacity for acquiring and refining motor skills, underpinning everything from mastering complex tasks like playing a musical instrument to everyday behavioural adjustments like walking on slippery surfaces. Traditionally, such learning has been conceptualized as an implicit process of the motor system, with neural studies often focused on the activity of brain areas like sensorimotor cortex, striatum, or cerebellum (Doya, 1999; Lalazar and Vaadia, 2008; Taylor et al., 2014). In recent years, however, multiple lines of behavioural evidence have demonstrated that motor learning can also be supported by the use of explicit cognitive processes (Mazzoni and Krakauer, 2006; Keisler and Shadmehr, 2010; Taylor and Ivry, 2011; Seidler et al., 2012; Galea et al., 2015; Holland et al., 2019). In classic visuomotor adaptation tasks, such as when the relationship between hand motion and visual feedback is altered during reaching movements, it has been shown that significant a proportion of subjects will adopt a re-aiming strategy during learning whereby they deliberately aim their hand in a direction that counteracts the cursor rotation (Taylor et al., 2014; de Brouwer et al., 2018). Notably, these studies indicate that such strategy use is highly variable across individuals, with those who exhibit a larger strategic contribution to learning tending to exhibit a faster reduction in visuomotor errors.

While the processes supporting implicit learning have been well characterized (e.g., by theories based on optimal feedback control, Scott, 2004), the component processes supporting explicit learning remain nebulously defined. Indeed, recent work has linked explicit learning to myriad cognitive processes such as working memory, attention, and inhibitory control, and has shown that individual differences in these processes relate to performance on motor tasks (Seidler et al., 2012; Galea et al., 2015). Outside the domain of the sensorimotor literature, neuropsychological studies discuss these processes under the umbrella term of ‘executive functions’ (EF; Karr et al., 2018), and indeed a large literature has linked motor coordination and control to the development of EF and general intelligence (Martin et al., 2010; Smits-Engelsman and Hill, 2012). Moreover, several studies have directly linked executive function to individual differences in whole-brain network structure (Reineberg et al., 2015; Reineberg and Banich, 2016). Much of this literature has specifically focused on the intrinsic functional organization of a set of higher-order cognitive regions comprising the default-mode, frontoparietal, and ventral-attention networks. In particular, an influential tripartite model proposed by Menon (2011) has linked various facets of higher-order cognition and related disease states to the integrity of functional interactions between this specific collection of networks.

Collectively, these findings offer a twofold prediction: Firstly, they predict that individual differences in motor learning performance can be explained by individual differences in EF. Secondly, they further predict that motor learning performance can be explained by individual differences in the intrinsic functional architecture of brain networks known to support EF. In order to explore both of these predictions, we tested whether individual differences in sensorimotor adaptation are associated with established behavioural and neural measures of general EF. In the first study, we had subjects perform both a standard visuomotor adaptation task and a task directly associated with executive functioning (a trail-making task), and found that the speed at which subjects reduced their error could be explained by their Trails performance. In the second study, we reanalyzed data from two of our own previously published sensorimotor adaptation studies (Areshenkoff et al., 2022, 2023), and found that consistent patterns of resting-state functional connectivity within tripartite cognitive brain networks were associated with more rapid learning in both tasks. By contrast, we found that motor learning performance could not be predicted based on the resting-state functional connectivity of sensorimotor-related brain networks.

## 1 Methods

### 1.1 Overview of tasks

We studied motor adaptation using a classic visuomotor rotation (VMR) task across three experiments: one behavioural, and two in an fMRI scanner (these latter two have been previously reported in Areshenkoff et al., 2022, 2023, see Table 1). In each task, subjects performed center-out reaching movements to launch a cursor towards a cued target in a circular array (see Figure 2; top left). After a set of baseline trials, a visuomotor rotation was abruptly introduced, such that the cursor travelled at a clockwise angle of 45*^◦^* relative to the subject’s hand movement. This required that subjects learn to adjust their hand movements in a counterclockwise direction to counteract the rotation of the cursor. In such tasks, the contribution of explicit processes to performance are generally greatest during the early stages of learning (Taylor et al., 2014), and we have previously found that changes in functional connectivity within and between cognitive brain networks are greatest during this early learning period (Areshenkoff et al., 2022). For these reasons, our present analyses focus on subjects’ performance error during this early learning period.

**Table 1:** Summary of the three experiments reported in this paper. Two of these, for which we collected fMRI data, have been previously reported elsewhere (Areshenkoff et al., 2022, 2023).

| Experiment | Platform | N | Citation | fMRI | Response device |
| --- | --- | --- | --- | --- | --- |
| VMR-TMT | mTurk | 75 | – | No | Computer mouse |
| VMR1 | In person | 36 | Areshenkoff et al. (2023) | Yes | Touchpad |
| VMR2 | In person | 32 | Areshenkoff et al. (2022) | Yes | Joystick |

In the first experiment (TMT-VMR), subjects (*N* = 74) were recruited through Mechanical Turk to perform a visuomotor rotation task, followed by a Trail-Making Task (TMT; see Figure 2, top right). The TMT comprised two task blocks, each requiring subjects to trace a path between labelled nodes as quickly and accurately as possible. In the first block (A), nodes were numbered, and subjects traced the path in numerical order. In the second (B), nodes were labelled with both numbers and letters, and subjects alternated between the two types of nodes (i.e. 1 *→ A →* 2 *→ B →…*). The total time taken to complete the second block is typically larger, and the excess time (the difference B-A) strongly correlates well with measures of executive processes (Sánchez-Cubillo et al., 2009; Salthouse, 2011). Following the suggestions of Salthouse (2011), we derived scores for each subject by regressing the time taken to complete block B onto the time taken to complete block A, and defining a subject *i*’s *residual score* to be the 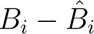, where 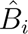 is subject *i*’s predicted block B time.

The VMR tasks in each of the VMR-TMT, VMR1, and VMR2 experiments were implemented nearly identically: Participants performed baseline trials in which they performed center-out target-directed movements. After these baseline trials, we applied a 45*^◦^* clockwise rotation to the viewed cursor, so that the cursor travelled at an angle relative to the movement of the subject’s hand/finger. At the beginning of the trial, a target (5 mm radius red circle) was shown on top of a grey ring with a radius of 60 mm (i.e., the target distance) centred around the start position. The target was presented at one of eight locations, separated by 45*^◦^* (0, 45, 90, 135, 180, 225, 270 and 315*^◦^*), in pseudorandomized bins of eight trials.

In each of the three tasks, subjects performed 64 baseline trials with no rotation, followed by a learning block of 160 trials (VMR1) or 200 trials (VMR and VMR-TMT). In the VMR1 experiment, the baseline and learning blocks were performed as part of separate scans, while in the VMR2 experiment they were performed consecutively as part of a single scan. Responses were executed in the TMT-VMR experiment using the computer mouse; in the VMR1 experiment with a touchpad; and in the VMR2 experiment with a joystick.

#### 1.1.1 Data quality screening

As has been frequently reported elsewhere (Rouse, 2015; Ahler et al., 2019; Chmielewski and Kucker, 2020), the quality of data collected through Mechanical Turk tends to be of a lower quality than the equivalent data collected in a laboratory setting. Ahler et al. (2019) found evidence that a substantial proportion of survey respondents (25-35%) show signs of significant data quality issues, such as deliberate “trolling” or inattentiveness, and that the noise introduced by these respondents can substantially attenuate estimated experimental effects. Chmielewski and Kucker (2020) reached similar conclusions, and found that, across several samples, a large proportion of respondents (roughly 40-60%) showed at least some evidence of questionable response validity. Given the focus on individual differences in our study, a particular concern of ours was that the inclusions of inattentive subjects, or subjects who otherwise failed to properly perform both tasks, might engineer the appearance of a relationship between TMT and VMR task performance. In order to guard against this possibility, we instituted a rigorous data quality screening procedure, which we describe below.

As our visuomotor rotation task was considerably more demanding than a simple survey, we wished to ensure that subjects exhibited behavior consistent with the performance we have observed in a laboratory setting. The end result was that, while a total of 253 subjects completed both the VMR and TMT tasks, we retained only *N* = 73 (30%) of these subjects after data filtering. As a clear justification of these removals, we illustrate the data cleaning pipeline in detail here.

Consistent with our previous work (Areshenkoff et al., 2022), we enforced a response deadline on individual VMR trials such that responses initiated more than 2 seconds after target appearance were considered invalid. As responses also required target discrimination, we also set a lower threshold of 100 msec in order to filter prepotent responses. We then removed subjects missing more than 25% of their total responses, as well as those missing more than 25% in either of the early learning (the first 34 learning trials) or late learning (last 36 trials), as these time windows were used to assess subjects’ total learning in the task. In total, 76 subjects were discarded in this stage.

We observed several subjects who appeared to ignore the target location entirely; responding either in the same direction on every trial, or in seemingly random directions. We interpreted this behavior as a sign of inattention, and quantified this tendency using the test statistic used in Rao’s uniformity test for circular variables (Rao, 1976). Loosely, a lower value for this quantity indicates that the subject’s errors are more uniformly distributed, and in practice performed well in identifying subjects’ whose responses concentrated around the target location, versus ignoring the target location entirely. For each subject, we computed this test statistic for the late learning period, when performance has generally reached asymptote. We call this statistic the subject’s *response concentration*.

When examining subjects’ total learning (their early and late errors), we observed what appeared to be a sub-cluster of subjects exhibiting high early error, as well as relatively weak learning (see Figure 1, top left). While this could be interpreted merely as poor learning, rather than inattention, we found also that these subjects tended to show more uniformly distributed errors (suggesting noisy responses, rather than merely incomplete learning). This noisy responding also appeared to be present during baseline.

**Figure 1:**
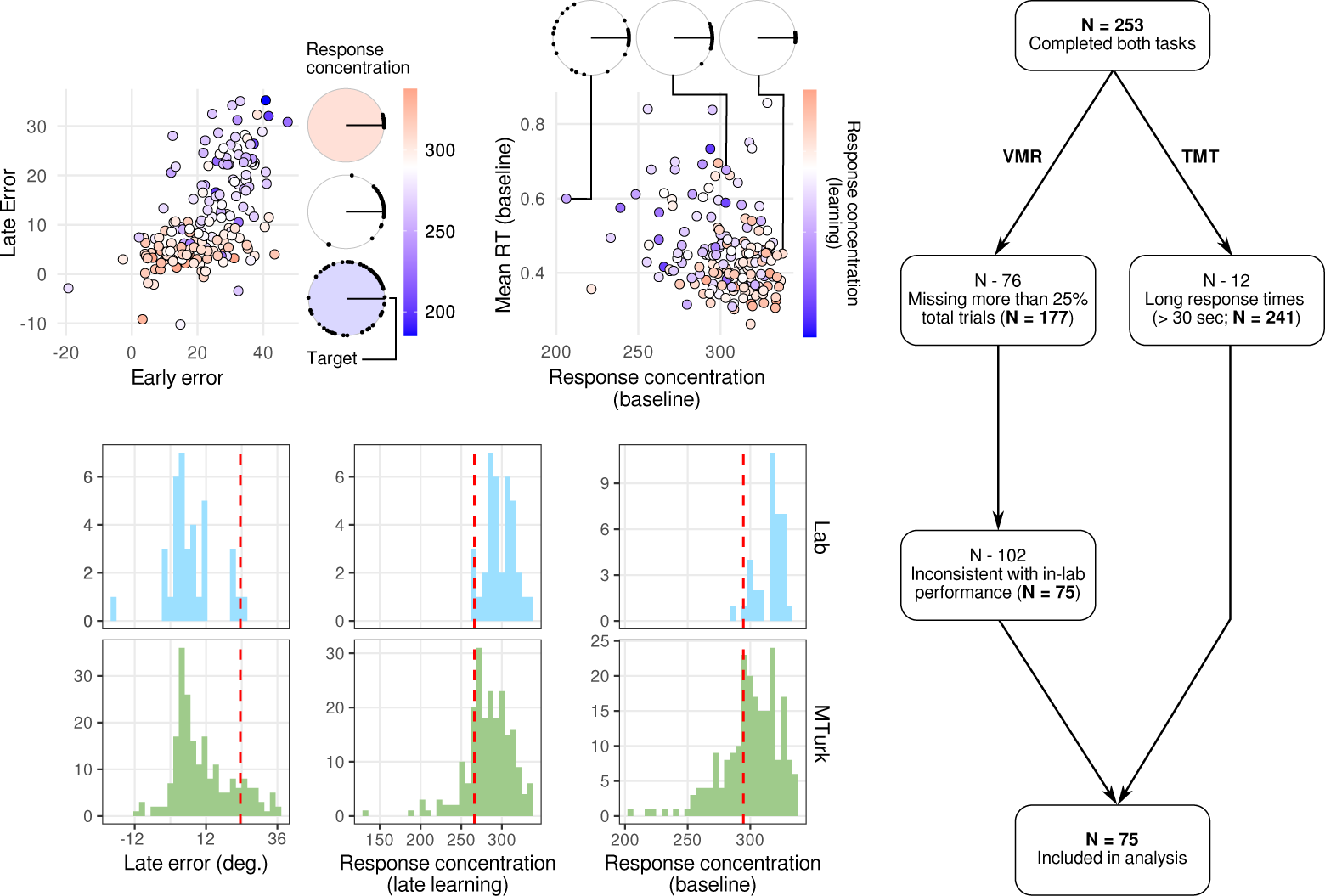
VMR performance for MTurk subjects. **Top left** panel depicts mean error during the early and late learning periods for all subjects who completed the VMR task. Color denotes *response concentration*, defined as the test statistic for Rao’s uniformity test (Rao, 1976). We identified what appeared to be a subgroup of subjects with high early error and poor total learning, who also consistently differed in their performance during baseline (e.g. high response variability). We used the behavioral data from the two in-lab VMR experiments in order to establish a normative baseline for performance, and excluded those subjects whose performance exceeded a 2.5% tail threshold on any of a set of three summary statistics (bottom panels). **Right)** Schematic diagram of MTurk data screening procedure. In total, *N* = 75 (29.6%) subjects were retained for analysis.

**Figure 2:**
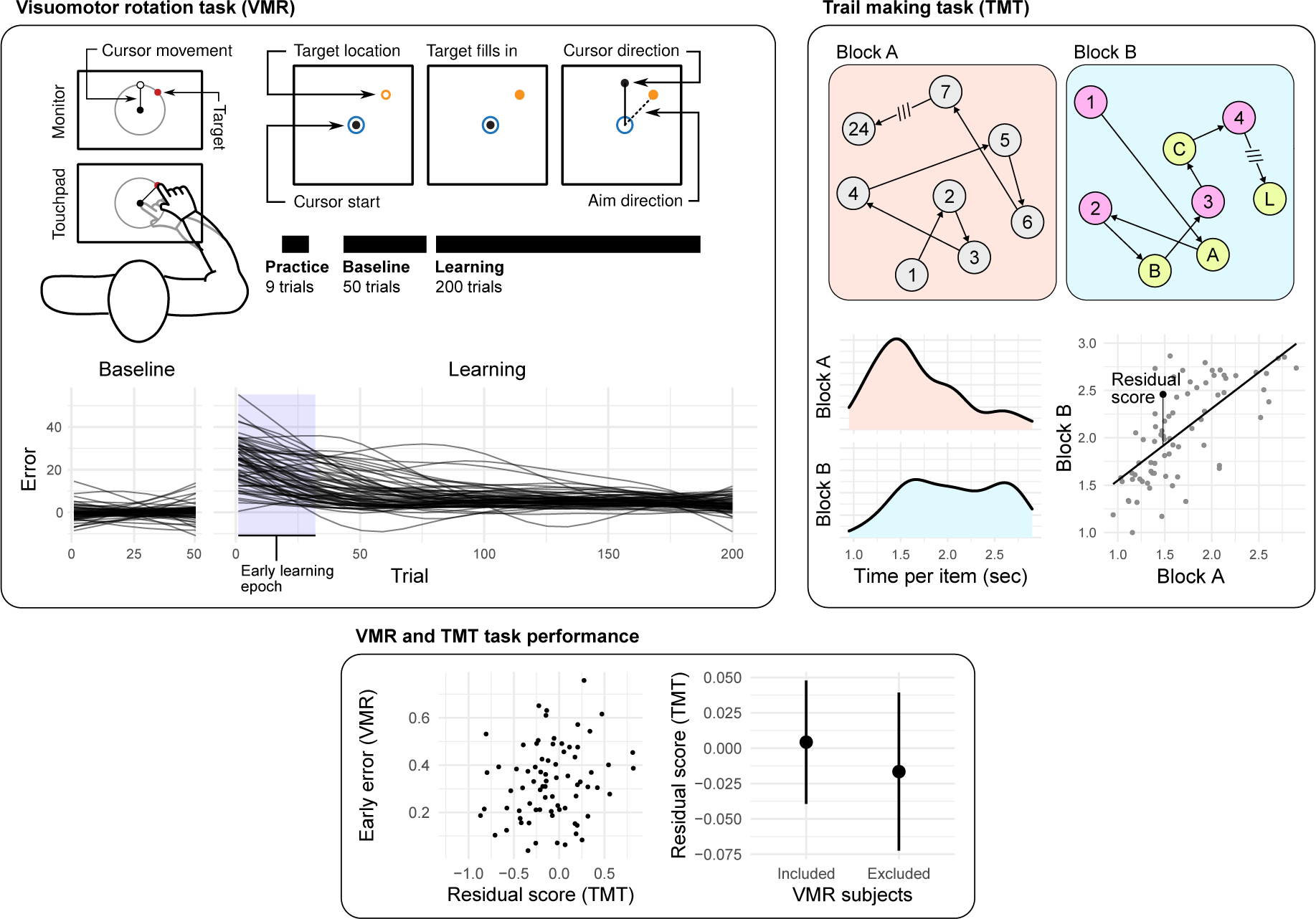
Visuomotor rotation task and subject behaviour. Online subjects performed both a trail making task (TMT) and a visuomotor rotation task (VMR). The TMT (**left**) comprised two blocks, each requiring subjects to trace a path between 24 labeled nodes. In the first block, nodes were labelled with numbers, and subjects were required to trace in numerical order. In the second, subjects alternated between letters and numbers (1 → *A* → 2 → *B* →…). In each block, we computed the mean time taken to complete each node, and regressed subjects’ reaction times in block B onto block A. The VMR task (**right**) required subjects to use a computer mouse to make centre-out reaching movements to a target located on a circular array.During the learning block, the correspondence between hand and cursor was rotated, so that the cursor moved at angle of 45*^◦^* relative to the hand. On each trial, we recorded the angular error between the final cursor position and the target, and (smoothed) subject learning curves are shown at the bottom right. Consistent with our previous analyses (Areshenkoff et al., 2022), we focus on the early learning period, in which individual differences in performance appear to be greatest.

In order to avoid imposing an arbitrary threshold on performance – which might be interpreted simply as removing subjects who did not “perform well” – we used the behavioral data from the two imaging tasks (VMR1 and VMR2), conducted in the lab, to establish a normative distribution of performance. We then excluded those MTurk subjects whose performance exceeded a tail-threshold (2.5%) of these distributions. Figure 1 (bottom left) shows the distributions of these variables for both the in-lab and MTurk versions of the VMR. It also clearly illustrates hints of bimodality in some of these variables, which might indicate a genuinely distinct subgroup of e.g. inattentive subjects, as opposed to mere poor-performers. In total, *N* = 75 subjects remained after this final exclusion step.

Performance in the trail-making task (TMT) was generally of higher quality, likely owing to the shorter and simpler nature of the task. We observed several subjects with extremely long response times for certain trials (i.e. several minutes), which we interpreted as distraction or inattention. In total, 12 subjects were excluded for response times greater than a liberal threshold of 30 seconds. We did not remove any other subjects on the basis of performance.

A simplified diagram of this screening procedure is shown in Figure 1 (right).

### 1.2 Behavioral data analysis

#### 1.2.1 TMT

Using the *N* = 75 subjects retained in both tasks, we computed mean reaction times for block A and B after Winsorizing with an upper tail threshold of 5% (Dixon, 1960). We then fit a standard linear regression model, regressing subjects’ mean block B reaction times block A, and used the residuals as subjects’ scores for analysis.

#### 1.2.2 VMR

Behavioral data from all three VMR tasks were analyzed identically. For all trials, we set upper and lower response time thresholds of 2 sec and 100 msec, respectively, and discarded responses that exceeded these bounds. For each trial, we then calculated the angular difference (the error) between the target and response location, and interpolated missing errors by fitting cubic splines to each subjects’ learning blocks. These splines were fit with knots at each observation, with smoothing penalty selected by generalized cross validation Craven and Wahba (1978).

For each subject, we defined the *early error* to be the circular mean error of the interpolated trials during the first four 8 trial blocks (comprising 34 trials; one presentation of each target location).

### 1.3 VMR1/VMR2 imaging procedures

We provide here an abridged description of the VMR1 and VMR2 data. More complete descriptions, and a thorough description of the neuroimaging preprocessing pipeline, can be found in Areshenkoff et al. (2022, 2023).

#### 1.3.1 VMR1

Forty-six right-handed subjects (27 females, aged 18-28 years, mean age: 20.3 years) participated in three separate testing sessions, each spaced approximately 1 week apart: the first, an MRI-training session, was followed by two subsequent MRI experimental sessions. One of these experimental sessions involved a task which is not analyzed here, and so we describe only the VMR testing session. Of these 46 subjects, 10 participants were excluded from the final analysis [1 subject was excluded due of excessive head motion in the MRI scanner (motion greater than 2 mm or 2° rotation within a single scan); 1 subject was excluded due interruption of the scan during the learning phase of the reward-based motor task; 5 subjects were excluded due to poor behavioural performance in one or both tasks (4 of these participants were excluded because >25% of trials were not completed within the maximum trial duration; and one because >20% of trials had missing data due to insufficient pressure of the fingertip on the MRI-compatible tablet); 3 subjects were excluded due to a failure to properly properly attend the second (unreported) task.

#### 1.3.2 VMR2

Forty right-handed individuals between the ages of 18 and 35 (M = 22.5, SD = 4.51; 13 males) participated in the study and received financial compensation for their time. Data from 8 participants were excluded due to either head motion in the MRI scanner (*N* = 4; motion greater than 2 mm or 2° rotation within a single scan) or their inability to learn the rotation task (*N* = 4), leaving 32 participants in the final analysis. Right-handedness was assessed with the Edinburgh handedness questionnaire (Oldfield, 1971). Participants’ written, informed consent was obtained before commencement of the experimental protocol. The Queen’s University Research Ethics Board approved the study and it was conducted in accordance with the principles outlined in the Canadian Tri-Council Policy Statement on Ethical Conduct for Research Involving Humans and the principles of the Declaration of Helsinki (1964).

### 1.4 Neuroimaging data analysis

#### 1.4.1 Covariance estimation and centering

Regions of interest were identified using the parcellation of Schaefer et al. (2018), and assigned network labels using the 17 network solution of Yeo et al. (2011). For each subject’s resting scan, we extracted standardized mean timecourses for ROIs belonging to the default mode (subnetworks A, B, and C), control (subnetworks A, B, C), and ventral attention (subnetworks A, and B). As part of a motor-network control analysis, we also extracted regions of the Somatomotor subnetwork A (comprising the hand area of the motor and somatosensory cortices), the Visual networks (A and B), and the dorsal attention network (A and B).

For each subject, we estimated covariance matrices using the linear shrinkage estimator of Ledoit and Wolf (2004). Because the resting state scans in the VMR1 and VMR2 experiments were of different lengths (297 and 177 imaging volumes, respectively), and because covariance estimates are biased (with the bias depending on the sample size), we worried that the difference in sample size may introduce systematic difference between tasks that would make cross-task prediction difficult. To remedy this, we centered the covariance matrices estimated from the VMR2 task so that the grand mean covariance matrices of each task aligned. The procedure we used is described in more detail in Areshenkoff et al. (2022), but we present here a basic account. All calculations are implemented in a freely available R package in a repository at https://github.com/areshenk-rpackages/spdm.

For each task, we estimated the mean covariance using the procedure outlined by Arnaudon et al. (2012); obtaining grand mean covariance matrices 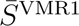 and 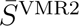. We then centered the VMR2 task to VMR1 by parallel transport. Specifically, for each covariance matrix 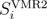 in VMR1, we projected it onto the tangent space at 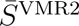, parallel transported the resulting tangent vector to 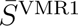, and projected it back onto the space of covariance matrices. While the underlying calculations are more complex, the conceptual meaning behind them is simply that the sample VMR2 is translated to have mean equal to VMR1.

#### 1.4.2 Predictive model

We used subjects’ resting state functional connectivity in the VMR1 and VMR2 experiments to predict their early error during subsequent task performance. The predictive model itself was trained on the data from the VMR1 task, and the VMR2 data was used in order to assess its out-of-sample predictive performance.

We fit our predictive model to the tangent vectors around the mean of VMR1, so that positive (resp. negative) values denote higher (resp. lower) than average functional connectivity at rest. We then vectorized each of these matrices as used the entries as features for our predictive model. Call the resulting matrix of predictors **X** = [**x_1_**, **x_2_***,…,* **x_n_**], where **x_i_** is a vector containing the vectorized resting state functional connectivity of the *i*’th subject.

We fit a ridge regression model to the VMR1 dataset with separate penalty terms for each network pair (e.g. DefaultA-ContlB) using the model described by (Van De Wiel et al., 2016), and implemented in the R package GRridge (van de Wiel and Novianti, 2022). These parameters were tuned using an internal leave-one-out cross-validation loop, so that networks whose regions contributed relatively little to overall predictive performance were shrunken more harshly relative to predictively useful networks. The precise arguments were as follows:

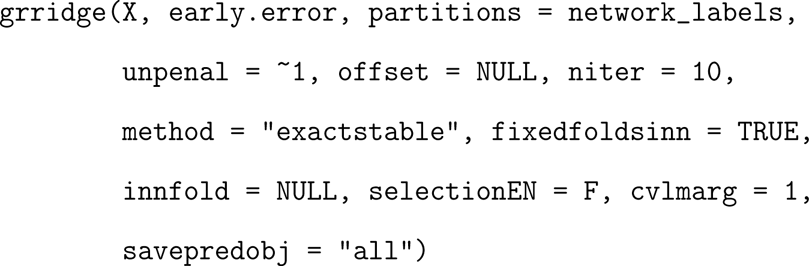

The resulting parameter vector *β* contains, for each entry in the covariance matrix, its estimated conditional contribution to subjects’ early error. Predictions for the VMR2 task were then derived using subjects’ tangent vector after centering to the mean of VMR1.

## 2 Results

### Trial-making performance predicts individual differences in sensorimotor adaptation

To test our first hypothesis that subjects’ executive function is a predictor of learning rate during sensorimotor adaptation, we examined the relationship between subjects’ performance on the TMT and VMR tasks in our mTurk data. Note that the mTurk data required substantial cleaning prior to analysis, and a large number of subjects were removed due to an apparent failure to properly attend to one or both tasks (see supplement for detailed exclusion criteria). Predictably, as the VMR task was considerably longer and more difficult, the majority of excluded subjects were removed due to unacceptable VMR performance (e.g. always responding in the same direction, too few valid trials within the early learning period). Our exclusion pipeline was highly aggressive, as we feared that inattentive subjects may perform poorly on both tasks, and thus give the appearance of a correlation between TMT and VMR task performance even if none exists. Still, we observed no significant difference in TMT performance between subjects who were included/excluded based on the VMR task (Figure 2, bottom; Welch’s test; *t*_197.64_ = 0.29411, *p* = 0.77, CI = [*−*0.120.16]). Consistent with our first hypothesis, in our retained subjects displaying acceptable performance in both tasks (*N* = 75), we observed a significant correlation between early error in the VMR task and residual score in the TMT (*r* = 0.25, *t*_73_ = 2.2054, *p* = 0.03), such that subjects who exhibited a better (lower) residual score in the TMT task displayed better (lower) early error during VMR learning. Together, these findings support the notion that individual differences in motor learning performance are associated with individual differences in executive function.

### 2.1 Baseline intrinsic cognitive network structure predicts individual differences in motor learning performance

Explicit, strategic contributions to sensorimotor adaptation performance have been shown to dominate the early stages of learning, and our previous work has linked individual differences in these explicit processes to the engagement of higher-order cognitive networks during learning, such as the default mode network (Areshenkoff et al., 2023). In turn, a substantial literature (noted in the Introduction) has linked the static, resting-state structure of such networks to individual differences in executive functions and general cognitive ability (Reineberg et al., 2015; Reineberg and Banich, 2016). Much of this literature has causally implicated regions of the VAN in the engagement of cognitive control in response to environmental demands (Menon and Uddin, 2010; Ham et al., 2013; Jiang et al., 2015; Parro et al., 2018), and has specifically linked functional interactions between the VAN and FPN/DMN networks with executive functions (Menon, 2011). Collectively, this large body of work predicts that individual differences in motor learning, which we have shown above reflects subject differences in executive function 2, may be further reflected in individual differences in resting-state cognitive network architecture.

To test this second hypothesis, we examined whether individual differences in resting-state functional connectivity between the DMN-FPN-VAN, which have been shown in previous work to be correlated with executive functioning (e.g. Reineberg et al., 2018), also exhibit patterns of correlation with VMR performance. To this end, we reanalyzed two of our previously reported datasets (Areshenkoff et al., 2022, 2023), in which subjects performed a standard VMR task in an MRI scanner. In each experiment, we collected resting-state fMRI scans prior to task performance, and we used these data to estimate intrinsic functional connectivity between regions of the DMN, FPN and VAN networks as defined by Yeo et al. (2011) and (Schaefer et al., 2018).

Using the data from Areshenkoff et al. (2023) (VMR1; N=36), we built a predictive model for subjects’ early error using the pairwise functional connectivity between regions of these three networks. Our approach was similar to our previous work in Areshenkoff et al. (2023), and utilized a sparse regression model with separate penalty terms estimated for coefficients associated with each pair of subnetworks. This allowed the model to select for connections between subnetwork pairs that were most strongly associated with learning performance, while controlling for the effect of other subnetwork pairs. A schematic diagram of this model is shown in Figure 4. We tuned the model through leave-one-subject-out cross-validation on the VMR1 dataset, and then used VMR2 as an independent test set in order to evaluate the model’s performance. Critically, this analysis revealed a significant positive correlation between the actual and model-predicted early error in the VMR2 sample (*r* = 0.4, *t*_30_ = 2.3943, *p* = 0.02, CI = [0.06, 0.66]). The fact that in the two datasets the separate groups of subjects used different response devices to perform the VMR task (summarized in Table 1), as well as different scanning parameters), indicates that motor learning performance can indeed be predicted from individual differences in resting-state cognitive functional network structure.

**Figure 3:**
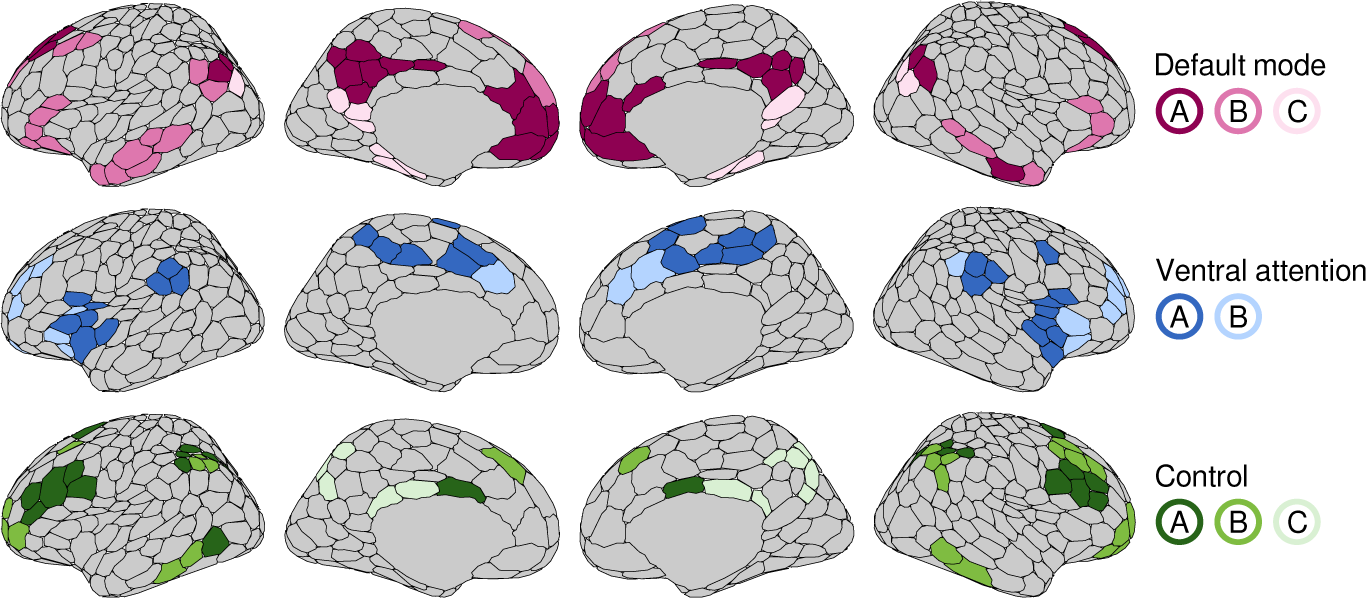
Network identification. We studied resting state functional connectivity between default mode, ventral attention, and control network. Regions of interest were extracted based on the parcellation of Schaefer et al. (2018), and assigned network labels based on the the 17 network solution of Yeo et al. (2011). For each subject in each task, we estimated correlation networks using the mean BOLD signal within each region.

**Figure 4:**
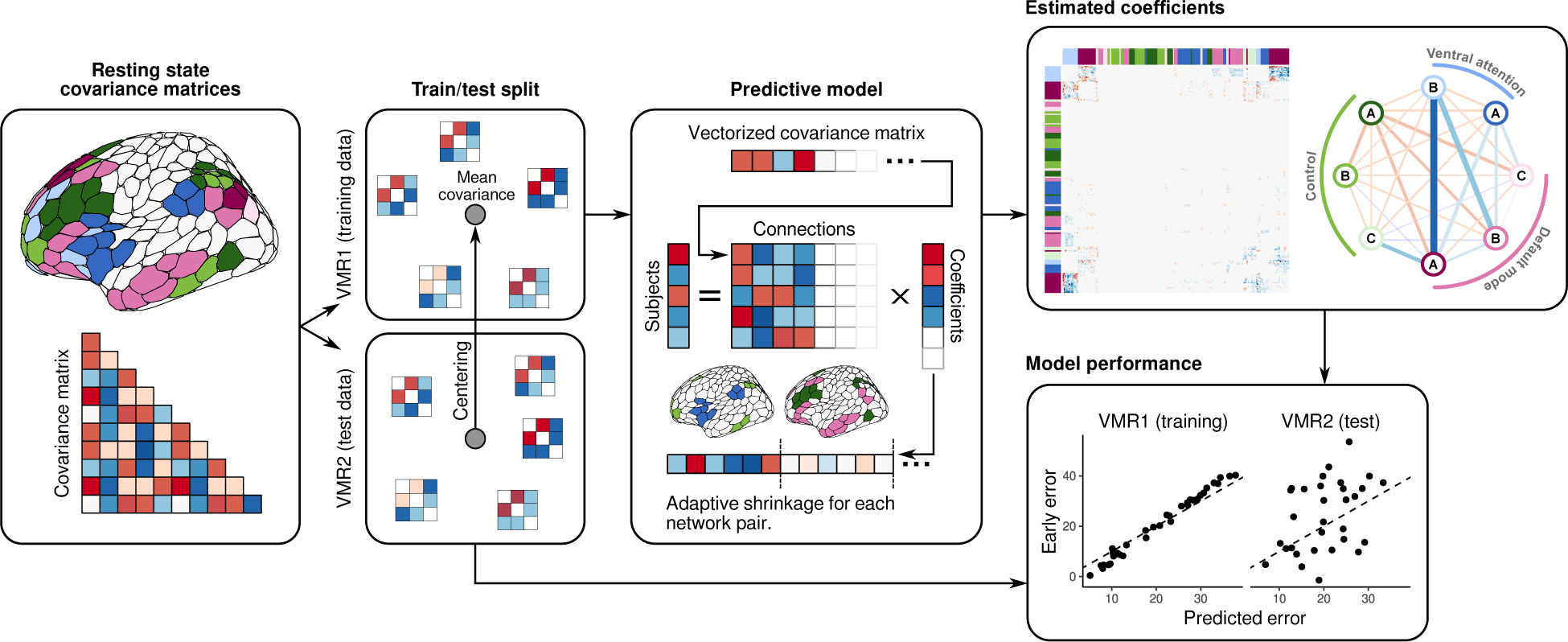
Predicting early learning performance resting-state functional connectivity. Using the VMR1 dataset as a training set, we trained a regularized linear model to predict subjects’ early error. We used the group-regularized ridge regression model of Van De Wiel et al. (2016), with separate penalty terms for connections between each pairs of subnetworks (e.g. DMNa-VANb, CNTb-DMNc). We evaluated our model using the the VMR2 data as a test set, the result of which are shown in the bottom right panel (Correlation between observed and predicted error: *r* = 0.4, *t*_30_ = 2.3943, *p* = 0.02, CI = [0.06, 0.66]). Model coefficients are displayed in the top right panel, along with mean coefficients for connections in each network-pair. The largest coefficients were seen between the ventral attention subnetwork B, and the default mode subnetworks A and B.

Given the success of this model, we next examined what specific subnetwork functional interactions were driving the prediction of motor learning performance. The bulk of the retained coefficients were associated with connections between regions of the ventral attention subnetwork B (the anterior portion of the ventral attention network, comprising the anterior insula and anterior-medial prefrontal cortex), and regions of the default mode subnetworks A and B (the core and dorsal medial subnetworks, respectively). The fitted model is shown in Figure 4 (top right).

On average, we found that greater resting-state connectivity between these pairwise networks was associated with a reduction in early visuomotor error. However, closer inspection of the VANb-DMNa coefficients (comprising the largest retained effects) revealed substantial heterogeneity in the connections between individual regions. In particular, we observed what appeared to be a clear laterality effect, in which increased connectivity between DMNa and regions of the right-hemisphere VANb were most strongly associated with lower error (see Figure 5). This effect is consistent with a large literature suggesting a hemispheric bias in the ventral attention network, in which the right anterior insula and cingulate cortex are thought to signal the engagement of both the default-mode and frontoparietal networks in order to meet environmental demands (Menon and Uddin, 2010).

**Figure 5:**
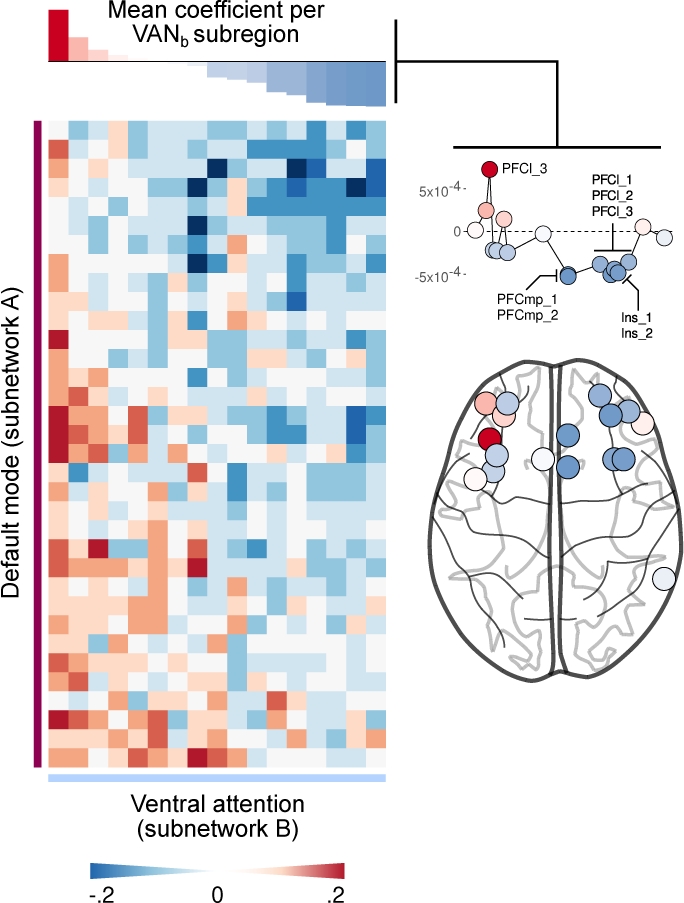
Laterality of VANb/DMNa coefficients. Coefficients for VANb/DMNa connections displayed an apparent lateralization, wherein stronger connections involving the right-hemisphere regions of the VAN were more strongly associated with lower early error.

### 2.2 Correlations between early visuomotor error and between-network functional connectivity

While our predictive model identified VANb-DMNa and VANb-DMNb connectivity as the dominant predictors of motor learning performance, these effects reflect partial correlations after controlling for the connectivity between other subnetworks. As raw correlations are more commonly reported in the related literature, we also computed the correlations between early visuomotor error and the average between-network functional connectivity of all subnetwork pairs (Figure 6). The center panel in this figure depicts, for each of the VMR1 and VMR2 tasks, the correlation between early error and between-network connectivity for each subnetwork pair. Note that, with a few exceptions, most networks pairs fall roughly along a unity line (with slope 1 and intercept 0), suggesting similar correlations between connectivity and performance across both tasks. This figure reveals that, on average, increased between-network functional connectivity for these higher-order cognitive was associated with a reduction in early error, with the strongest negative correlations with performance typically involving connectivity between the default-mode and ventral-attention networks.

**Figure 6:**
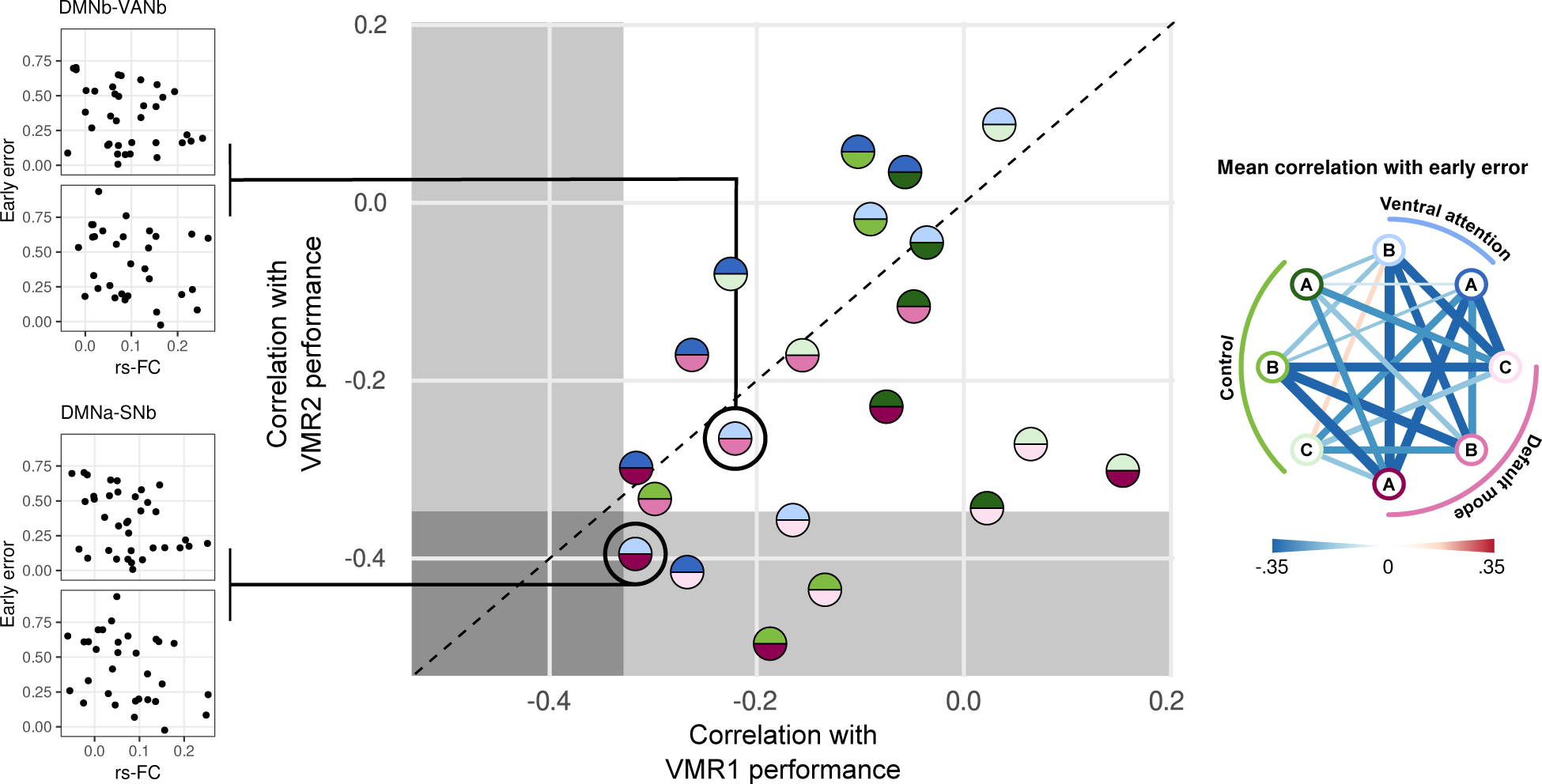
Between-network functional connectivity during rest and early learning. For each of VMR1 and VMR2, we computed the mean resting-state functional connectivity between subnetworks of the default-mode (DMN), control (CNT), and ventral attention networks (VAN). These correlations are depicted in the center panel (shaded regions indicate critical regions for two-tailed t-test). The leftmost panels depict early error and mean rs-FC for DMNb-VANa and DMNa-VANb connections, which were the selected for in our model as the most predictively important connections. The rightmost network depicts the mean correlation with early error for each pair of functional networks.

### 2.3 Control analyses

One possible alternative explanation for our effects, which we interpret as reflecting the importance of higher-order executive processes and their associated brain networks in motor learning performance, is that they instead reflect a non-specific relationship between resting network structure and performance; e.g., a global effect related to whole-brain functional connectivity, or to general fatigue at the time of testing. To rule out these alternative explanations, we first examined whether we could replicate our across-task prediction of early error using sensory-motor, rather than cognitive, brain networks. In the domain of motor sequence learning, prior work (Mattar et al., 2018) has shown that baseline, resting-state functional connectivity between visual and motor cortical regions is correlated with subjects’ subsequent learning rates, suggesting that the brain’s baseline sensorimotor functional architecture can predict future learning performance. To test a similar idea in the context of sensorimotor adaptation, we performed the same analysis done in Fig 4 but for regions in the primary somatomotor and visual networks, as well as the dorsal attention network (DAN). Together, these networks comprise the cortical circuitry supporting the execution of the motor response, and the sensory-error based updating of an internal model which it believed to be responsible for the bulk of implicit adaptation in the VMR task (Krakauer, 2009). Critically, as shown in Figure 7 (left), using these sensorimotor networks we observed no ability to predict VMR2 task performance based on a model fit of the data from VMR1 (*r* = *−*0.06, *t*_30_ = *−*0.34, *p* = 0.74).

**Figure 7:**
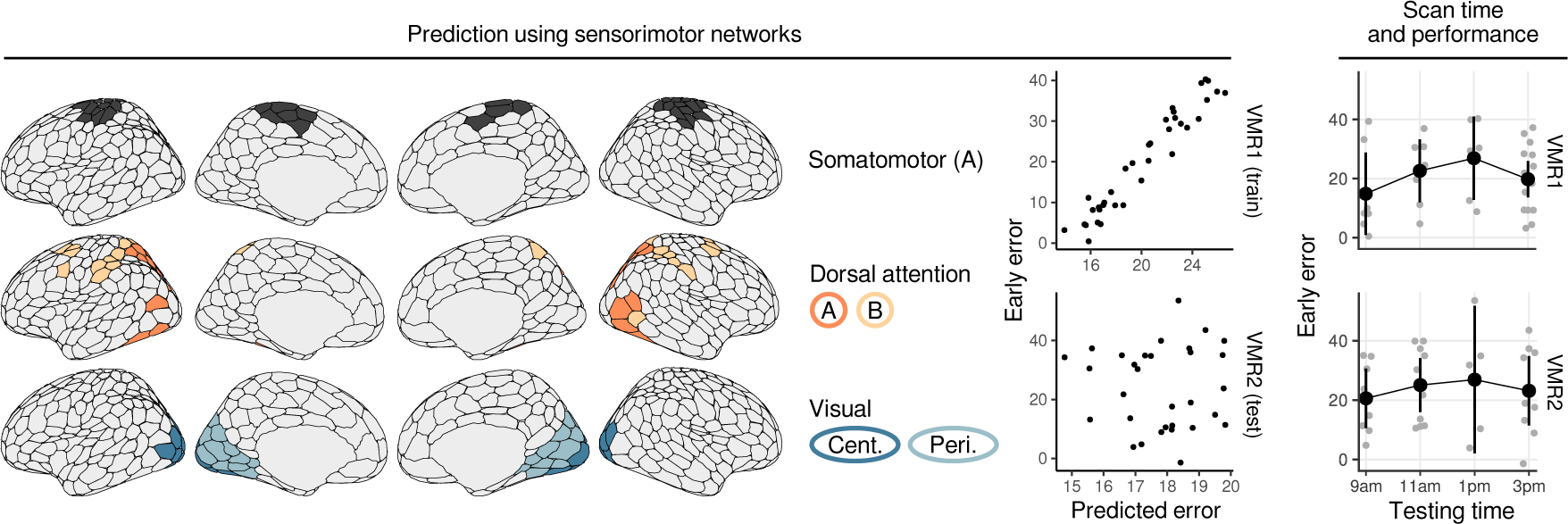
Control analyses. We fit a second, identical predictive model using connectivity between regions of primary motor and visual networks, and the dorsal attention network (left). This model failed to successfully predict subjects’ early error in the separate, VMR2 task. We also examined the relationship between subjects’ performance (early error) in the VMR1/VMR2 tasks, and the time of their testing session, in order to rule out any general effects of fatigue. Here, as well, we found no effect on subjects’ behavior.

Another alternative explanation, given that testing times occurred throughout the day (from 9am to 5pm), is that differences in fatigue/circadian rhythm between early morning and afternoon testing sessions may have caused differences in both sensorimotor performance and resting state functional connectivity. To test this, we directly examined the relationship between VMR performance (early error) in each of the imaging tasks, and the time at which the subject was scanned (see Figure 7, right, for mean error per time slot). Note that we did not observe any apparent differences in performance in either VMR task based on time of day, nor did a one-way ANOVA reveal a significant effect of testing time on early error in either task (**VMR1**: *F*_3_,_31_ = 1.091, *p* = 0.37; **VMR2**: *F*_3_,_29_ = 0.24, *p* = 0.87). Taken together, the above findings suggest that the relationship between cognitive network architecture and sensorimotor adaptation performance is not due to extraneous factors, but instead relates to the specific cognitive functions performed by those networks.

## 3 Discussion

Traditionally, studies of motor learning have focused on how neural computations within the motor system drive adaptation. However, recent work has demonstrated that cognitive strategies, which encompass myriad processes such as working memory, attention, and inhibitory control, can also play an important role in learning and in fact, explain individual differences in performance. Given that, outside the domain of motor learning, these myriad processes reflect constituent components of executive function, here we tested the dual proposition that (1) behavioural measures of executive function and (2) spontaneous activity within brain networks linked to executive function, would both predict individual differences in motor learning performance. Consistent with these predictions, we found that performance on the trail-making task, a classic measure of executive function, predicted the speed of visuomotor adaptation in an online sample of participants. In addition, using discovery and replication samples of fMRI data, we found that pre-learning resting-state functional interactions between the VAN and DMN, networks implicated in executive function, were also predictive of visuomotor adaptation performance. Together, these findings underscore the importance of considering individual differences in broader cognitive abilities and their associated brain networks when investigating the variability in basic motor learning processes. Prior work has emphasized two contrasting views regarding the nature of functional brain network organization. One of these has focused on the flexible, context-dependent, and “state”-like features of brain networks, wherein they dynamically reconfigure in response to tasks, moods, or even fleeting thoughts (Finn et al., 2015; Geerligs et al., 2015; Gordon et al., 2017a, 2022, 2017b). A second view has emphasized the stable, subject-specific structure of these networks; generally suggested to reflect the underlying, white-matter structure (Laumann et al., 2015; Mueller et al., 2013) governed by genetics and long-term patterns of activity. As shown by Gratton et al. (2018), intrinsic resting-state functional brain structure is stable over time, and highly individual-specific. Moreover, these static subject differences appear to dwarf any changes in activity associated with experimental conditions, temporary cognitive states or daily fluctuations, a finding we have replicated in several of our own analyses, in both Humans and non-Human primates (Areshenkoff et al., 2021, 2022, 2023). These findings have often been used to justify the use of baseline (’resting-state) subtraction in order to isolate task-related changes in connectivity Gratton et al. (2018), but our results here – that individual differences in motor learning performance can be predicted by pre-existing, trait-like patterns of functional connectivity within cognitive control networks – suggest that such procedures might also discard important subject-specific information contributing to learning.

These stable differences in brain network structure have in turn been linked to individual differences in general cognitive functioning (Wang et al., 2011; Finn et al., 2015). Note that the neural substrates of executive functions are difficult to relate to the activity of individual brain regions. Rather, the consensus is that executive processes rely on the integrity of large-scale brain networks, comprising multiple regions which coordinate in order to facilitate adaptive behaviour (Bettcher et al., 2016; Santarnecchi et al., 2021). In particular, Menon (2011) has linked various facets of cognition and disease to the integrity of the cortical functional activity spanned by regions of the DMN, FPN and VAN. At a high level, the VAN appears to be important for the detection of behaviorally relevant stimuli (Corbetta and Shulman, 2002), and in regulating the relative engagement of the FPN and DMN in order to facilitate either externally directed or introspective cognition. The proper functioning of this triple-network architecture is thought to underlie the ability of an individual to flexibly adapt to changing environmental demands, and dysfunction within and between these networks appears to underlie a large number of neuropsychological and psychiatric conditions (Menon, 2011; Shaw et al., 2021). Moreover, their functional integrity has been linked to individual differences in executive functions (Reineberg et al., 2015; Reineberg and Banich, 2016).

Individual regions of the VAN – especially the right anterior insula and cingulate cortex – have been causally implicated in signalling the engagement of cognitive control in response to environmental demands (Menon and Uddin, 2010; Ham et al., 2013; Jiang et al., 2015; Parro et al., 2018). Functional connectivity between the VAN and the DMN/FPN in turn appears to correlate with changes in cognition and memory during aging (La Corte et al., 2016), and is altered after traumatic brain injury (Lu et al., 2022). These accounts are consistent with a large body of imaging and neural recording work, both in Humans and non-Human primates, causally implicating parts of the insula and cingulate cortex in signaling the need for ongoing cognitive control (Grohn et al., 2024). Note also that we have previously implicated regions of the DMN, in particular, in explicit motor learning processes (Gale et al., 2022; Areshenkoff et al., 2023), which is itself consistent with recent work suggesting that the DMN is important for tasks requiring responding based on internally-generated rulesets, or episodic memory of previous trials (Smallwood et al., 2021).

Given these findings, and other work linking the development of motor coordination to executive functions (Martin et al., 2010; Smits-Engelsman and Hill, 2012), we suggest that research focusing solely on the computations performed by the motor system during sensorimotor adaptation (e.g. work which attributes adaptation primarily to the updating of an internal forward model), neglects the critical cognitive machinery that underlies a substantial degree of individual variability in motor learning performance. In turn, this machinery appears to be supported, in large part, by the general integrity of whole-brain cognitive networks, rather than by systems engaged solely during the learning process itself.

### 3.1 Future directions

Although we discuss our findings in terms of executive functions (EF) and general cognitive ability, our assessment of subjects’ EF was indirect. Our measure of subjects’ trails performance has been shown to correlate faithfully with general EF (Salthouse, 2011), but a more comprehensive task battery would allow the assessment of exactly which general or specific executive subcomponents were most strongly associated with motor learning performance. The patterns of resting-state functional connectivity which we have found correlate with learning have also been related to general cognitive ability, although a more comprehensive task battery would likewise allow researchers to identify patterns of connectivity specifically associated with different components of EF. Thus, such an approach could shed more light on the specific processes underlying individual differences in motor learning.

An important open question, which is the focus of much of our previous work (Areshenkoff et al., 2022, 2023), concerns the cognitive and neural substrates of explicit motor learning processes (i.e. those involving cognitive strategies), as opposed to implicit adaptation. Our previous work has specifically implicated regions of the frontoparietal and default mode networks in these explicit learning processes, which is itself consistent with work implicating these systems in conscious monitoring during motor learning (e.g. Slachevsky et al., 2003). A more comprehensive assessments of subjects’ executive functions, along with an assay of explicit and implicit processes during the initial learning period (e.g. as in Taylor et al., 2014) would certainly be extremely informative regarding the processes supporting these competing learning systems.

Finally, our findings predicting individual learning ability have implications for both neurorehabilitation (for aging, injured, or diseased populations) and neuroeducation (for children or older trainees). Unlike predictors based on task performance, resting-state scans offer a practical advantage as they can be applied to individuals unable to perform tasks or remain still during task-based imaging sessions. As such, our ability to identify stable trait-like subject factors that predict sensorimotor performance could be useful in patient selection for clinicial trials or optimizing future learning outcomes in rehabilitation or training settings.

## Acknowledgement

This work was supported by operating grants from the Canadian Institutes of Health Research (CIHR) awarded to J.P.G. (MOP126158). J.P.G. was also supported by an NSERC Discovery Grant, as well as funding from the Canadian Foundation for Innovation. J.R.F was supported by a Natural Sciences and Engineering Research Council of Canada research grant (05944-2019), and a Canadian Institutes of Health Research grant (156173). C.N.A. was supported by a Natural Sciences and Engineering Research Council (NSERC) graduate award. We thank Martin York and Don Brien for technical assistance.

